# *Enterocytozoon hepatopenaei* Proliferate in *Procambarus clarkii*: A Warning for Crayfish and Shrimp Aquaculture

**DOI:** 10.1101/2023.10.15.561606

**Authors:** Bingxin Ling, Yujiao Wu, Qing Yu, Chunxia Wang, Mengjiao Hu, Xianzhi Meng, Mengxian Long, Guoqing Pan, Zhonghuai Xiang, Zeyang Zhou, Jie Chen

## Abstract

The pacific whiteleg shrimp (*Litopenaeus vannamei*) and the crayfish (*Procambarus clarkii*) are the most productive aquatic animals in the world. The prevalence of the microsporidium *Enterocytozoon hepatopenaei* (EHP), an intracellular spore-forming unicellular parasite which leading retarded growth of *L. vannamei*, has caused severe economic losses in most shrimp farming country. In this study, we found that the wild *P. clarkii*, living in shrimp ponds with EHP outbreak, excreted white faeces after temporary laboratory culture. The hepatopancreas (HP) of symptomatic crayfish exhibited a lighter color and severely atrophied. H&E-stain showed tissue lesions in both hepatopancreas and intestine, and clustered microsporidian spores were filled in the cytoplasm of the cells. PCR using *EHPptp2* and two microsporidian-universal primers sets demonstrated the existence of EHP in the hepatopancreas, intestine and the white feces of *P. clarkii*. The EHP loads of 10^3^-10^4^ copies of *EHPptp2*/50 ng HPgDNA were detected by qPCR. Developing stages and mature spores of EHP were observed in hepatopancreas of *P. clarkii* through indirect-immunofluorescence assay (IFA) and transmission electron microscopy (TEM). Considering the large-scale cultivation of *L. vannamei* and *P. clarkii*, overlap farming areas between these two species as well as the ability of crayfish to crawl on land and water, our finding indicates the potential role of *P. clarkii* in the transmission of EHP, and it is an early warning for crayfish and shrimp farming.

## 1. Introduction

Pacific whiteleg shrimp (*Litopenaeus vannamei*) is one of the most productive aquatic animal in the world, reaching 5.8 million tons in 2020 (FAO, 2022). With the rapid expansion, disease problems plague the healthy development of shrimp farming. *Enterocytozoon hepatopenaei*, the pathogen of hepatopancreas microsporidiosis (HPM), is the microsporidium that can infect shrimp, resulting growth retardation and susceptibility to multiple infections with bacteria and viruses (Chayaburakul et al., 2004; Flegel, 2012). Since the first report of EHP in black tiger shrimp (*Penaeus monodon*) with stunted growth in Thailand (Tourtip et al., 2009), the pathogen has had a wider impact on *L. vannamei*, leading huge economic losses in most shrimp farming country and region (Ha et al., 2010; Wang et al., 2013; Tang et al., 2015; Tang et al., 2016; Biju et al., 2016; Shen et al., 2017; Tang et al., 2017; Kim et al., 2021). Aquaculture farmer lost their investment of manpower, resources, financial and time due to the chronic infection caused by EHP. For instance, Thailand’s national economy lost has rose to US$180 million from 2010 to 2016, and to US$232 million in 2018 (Shinn et al., 2018; Patil et al., 2021). India lost US$567.64 million due to EHP in 2018-2019 (Patil et al., 2021). Moreover, lost is even immeasurable in China (Wang et al., 2013). The spores of EHP are oval profile, and 0.7 × 1.1 μm in size (Tourtip et al., 2009). EHP has spore wall enveloping sporoplasm, in which it contains a nucleus, 5-6 coils of the polar filament, a posterior vacuole, polaroplast, an anchoring disk attached to the polar filament and organelles such as ribosome, endoplasmic reticulum, Golgi apparatus, and mitochondrial remnants named mitosome (Tourtip et al., 2009; Weiss and Becnel, 2014). The life cycle of EHP can be divide into proliferative stage, sporogonic stage, and infective stage (mature spores) (Tourtip et al., 2009; Vavra and Lukes, 2013; Chaijarasphong et al., 2021). At proliferative stage, cells divide firstly by binary fission, followed by multiple nuclear divisions without cell division (Ning, 2020). At sporogonic stage (sporogonic plasmodium), the electron-dense disk (EDD) forms the polar filament, and then dense secretions thickens the plasma membrane, accompanying with the sporoblasts beginning to form. Spore wall of the sporoblasts gradually thicken and spores develop into mature ones (Ning, 2020). EHP can be transmitted horizontally though water (Salachan et al., 2017), feces (Singh and Singh, 2018) and carcasses (Tangprasittipap et al., 2013; Jang et al., 2022). Besides *L. vannamei* and *P. monodon*, EHP was found proliferate in blue shrimp (*Penaeus stylirostris*) (Tang et al., 2015), giant river prawn (*Macrobrachium rosenbergi*) (Wang et al., 2022) and dragonfly (*Anax parthenope*, *Pantala flavescens*, and *Ischnura senegalensis*) (Kumar Dewangan et al., 2023). EHP can also be transmitted between live foods and other animals living in the same waters, such as polychaetes (Desrina et al., 2020), brine shrimp (*Artemia salina*) (Karthikeyan and Sudhakaran, 2020), false mussels (*Mytilopsis leucophaeata*) (Munkongwongsiri et al., 2022), marine crabs (Mani et al., 2022) and some macrofauna (Wan Sajiri et al., 2023). Although a variety of methods including specific pathogen free (SPF) shrimp larvae breeding has been taken, the outbreak of EHP remained frequently happening, suggesting a complex biological transmission pathway of EHP (Chayaburakul et al., 2004; Jang et al., 2022).

The crayfish (*Procambarus clarkii*), native to North America, is farmed all over the world and forms large wild populations now. It was introduced into China in the 1930s, then its production has soared yearly and reached 2.6 million tons in 2021 (Westman, 2002; Guo et al., 2017; Fisheries administration of the Ministry of Agriculture and Rural Affairs of the Peoplès Republic of China, 2022). Its main domestic breeding areas in China are concentrated in the middle and lower reaches of the Yangtze River at present (Guo et al., 2017; Fisheries administration of the Ministry of Agriculture and Rural Affairs of the Peoplès Republic of China, 2022).The crayfish and shrimp farming areas are usually overlapped in China, even plenty of shrimp ponds are converted from crayfish ponds. The crayfish is capable of crawling, digging and breeding, and can survive cold winters in burrows. They have the habit of climbing ashore, cross dikes and enter other bodies of water (Xie et al., 2008). These characteristics allow crayfish to transmit pathogens easily between different water bodies.

In this study, we found the wild crayfish *P. clarkii* which inhabited in the farming ponds of *L. vannamei* with slow growth syndrome were infected with EHP. The *P. clarkii* shed white faeces during temporary laboratory culture. Molecular detection, histopathological observation and ultrastructure analysis were used to demonstrate the present and proliferation of EHP in the hepatopancreas of *P. clarkii*.

## 2. Material and methods

### 2.1 Sample collection

The wild crayfish ranging from 9.5 to 10.5 cm were collected from a *L. vannamei* shrimp pond which suffered an outbreak of EHP in Chongqing, China. These crayfish were temporary reared in a 100 L ton within 40 L water at 26. The excrement and food residue were cleaned up, and half of the water was replaced by fresh ones every day. White feces were collected after one day of laboratory culture. Segments of white feces were fixed in 4% paraformaldehyde, while the remained were stored at -20. The hepatopancreas and intestines of healthy and diseased crayfish were sampled for EHP detection and histological analysis.

### 2.2 DNA extraction

Sampled hepatopancreas, intestine and white feces of the infected crayfish were grounded thoroughly. Genomic DNA was extracted by 2% CTAB (1.40 M NaCl, 0.55 M CTAB, 125 mM EDTA, 40 mM Tri-HCl, and 0.2%-1% β-Mercaptoethanol, pH=8.0) and 1 mg/ml protease K, and incubated at 65 °C for 2 h. After incubation, the equivalent value of phenol/chloroform/isoamylalcohol extraction was added to the digests. The extracted DNA in aqueous phase was successively precipitated by isopropanol and washed by gradient ethanol (90% and 70%). The purified DNA was dissolved in sterilized ddH_2_O, diluted to 50 ng/µL, and stored at -20°C.

### 2.3 EHP detection and the crayfish identification

The primer sets targeting universal for microsporidia and EHP polar tube protein 2 (*EHPptp2*) were used to identify the infection of EHP (Zhu et al., 1993; Baker et al., 1995; Wang et al., 2021; Wu et al., 2022). Two pairs of primers amplifying mitochondrial genes were employed to identify the crayfish (Folmer et al., 1994; Li et al., 2015). The details of primers used in this study were listed in Table 1. The PCR products were cloned into pESI-Blunt vector and sequenced (Sangon, China). Sequences of *SSU rRNA* (GenBank: MNPJ01000021.1), *EHPptp2* (GenBank: MT249228.1), mitochondrial cytochrome oxidase *c* subunit I (*cox*) (GenBank: JN000903.1) and mitochondrial control region (Displacement loop region) (D-loop) (GeneBank: KC556829.1) were download from NCBI. Sequence analysis was carried out using MEGA (11.0.13) (Tamura et al., 2021) and GeneDoc (Nicholas and Nicholas, 1997). qPCR was performed using Hieff qPCR SYBR® Green Master Mix (Yeasen, China) in LightCycler® 96 real-time PCR instrument (Roche, China), followed the instructions.

**TABLE 1.**
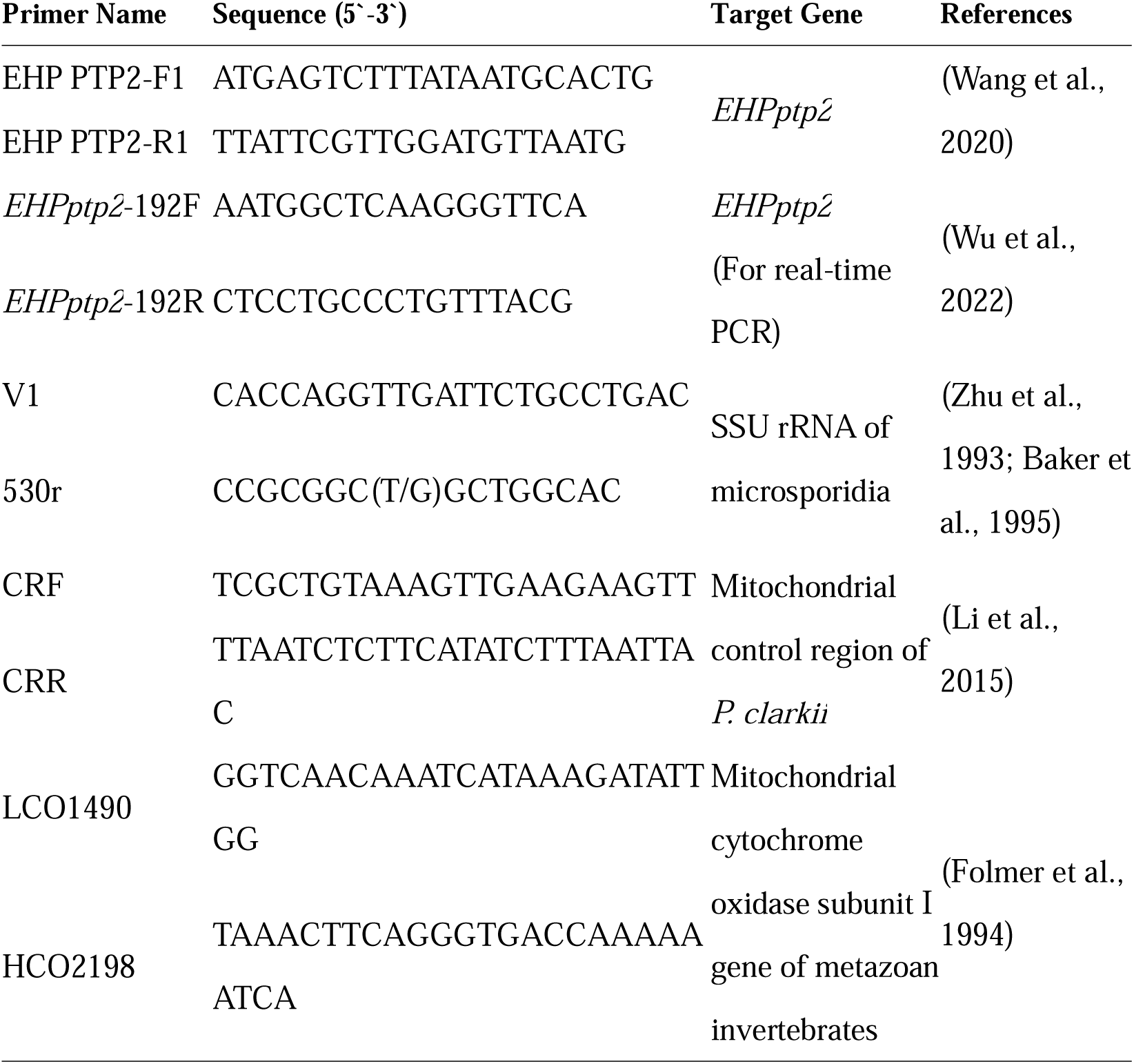
Primers used in this study.

### 2.4 Fluorescence microscopy and load assay of EHP

Tissue samples were homogenized and appropriate amount were used to detect the EHP spores through fluorescence microscopy. Samples were incubated with Fluorescent Brightener 28 (Sigma Aldrich, USA) and Propidium iodide (Thermo Fisher Scientific, USA) for 5 min, and then observed by fluorescence microscope (Olympus, Japan) using differential interference contrast and ultra violet filter (Zhao et al., 2020; Chen et al., 2021).

### 2.5 Histological analysis by H&E-staining

An appropriate tissue block was immediately cut from the live crayfish and fixed in Davidson’s AFA fixative. The tissues were embedded in paraffin and sliced to 5 µm thick. The paraffin sections were stained by the hematoxylin eosin staining kit (Beyotime, China), then stained with Fluorescent Brightener 28 (Sigma Aldrich, USA) for 10 minutes, and observed with microscope (Olympus, Japan) using differential interference contrast and ultra violet filter.

### 2.6 Indirect-immunofluorescence assay (IFA)

IFA was performed to specifically recognize different phases of EHP in paraffin sections prepared previously. Dewaxed paraffin sections were dealt with Citrate antigen repair solution (Beyotime, China) for 30 minutes at 98. After blocking, sections were incubated with anti-EHP antisera or unimmunized antiserum (negative serum) (diluent in 1:200), which is prepared by total proteins of EHP mature spores, at room temperature for 2 hours. Then the sections were incubated with Alexa Flour® 488 conjugate goat anti-mouse IgG (Thermo Fisher Scientific, USA) following washing steps. Fluorescent Brightener 28 (FB 28) and Propidium iodide (PI) were used to stain chitin layer of spores and nuclei of cells respectively for 15 min. All samples were sealed with ProLong® Gold antifade reagents (Thermo Fisher Scientific, USA). Imagines were carried out by Olympus FV1200 laser scanning confocal microscope (Olympus, Japan).

### 2.7 Transmission Electron Microscopy (TEM)

TEM was performed as described by Meng at el. (Meng et al., 2018) with slight modifications. Hepatopancreas and feces were cut into small pieces (≤1 mm^3^) and fixed in 5% glutaraldehyde over 24 hours. After washed with 0.1 M PBS buffer (pH=7.4) three times (10 minutes each), samples were postfixed in 1% osmium tetroxide for 2 h. The fixed tissues were embedded in epoxy resin Epon-812 and polymerized at 37 for 24 h and at 60 for 48 h. Ultrathin sections (60-90 nm) were collected on copper grids and stained with uranyl acetate and lead citrate. Micrographs were taken using the transmission electron microscope (HITACHI, Japan).

## 3. Results

### 3.1 *P. clarkii* in *L. vannamei* farming ponds suffered EHP outbreak was positive for EHP and exhibited white feces

The wild crayfish, collected from a *L. vannamei* farming ponds suffered an EHP outbreak, excreted white feces in the first five days under laboratory culture. Species identification, using primers set LCO1490 and HCO2198 (Folmer et al., 1994), CRF and CRR (Li et al., 2015) indicated the crayfish is *Procambarus clarkii* (Fig. S1 & Fig. S2). The hepatopancreas of the symptomatic crayfish was discolored and atrophied. Besides lightened color, part of the intestine exhibited empty (Fig. 1 A-C). Fluorescence microscopy of the tissues smear showed numerous microsporidian spores and germinated spore coats presented in hepatopancreas, intestine and white feces (Fig. 2). Genomic DNA isolated from these samples was subjected to PCR amplification using primers targeting *SSU rRNA* of microsporidia and *EHPptp2* (Fig. 3 A). These sequenced fragments were matched with the *SSU rRNA* and the conference *EHPptp2* respectively (Wiredu Boakye et al., 2017) (Fig. 3 C & D), demonstrating that the microsporidium found in wild *P. clarkii* is coincident with the conference EHP strain TH1. *EHPptp2* qPCR results showed 1.62×10^3^, 1.93×10^3^ and 5.61×10^4^ copies/50 ng gDNA loads in intestine, hepatopancreas and white feces of EHP infected *P. clarkii* respectively (Fig. 3 B).

**Fig. 1.**
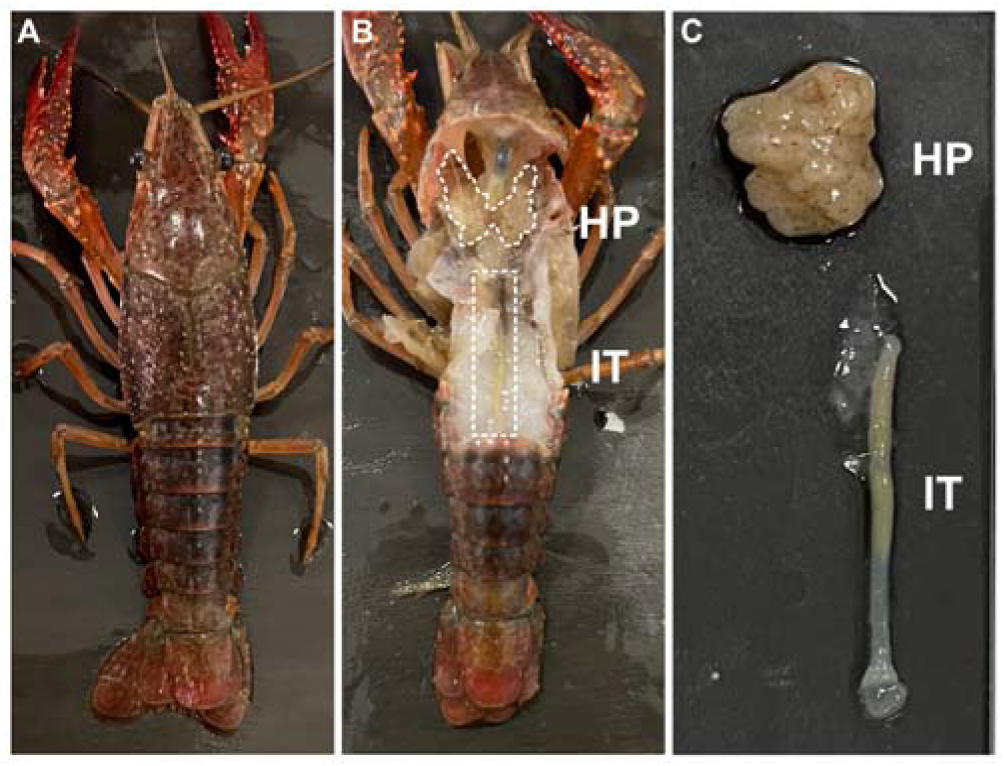
The anatomical morphology of digestive tract in infected *P. clarkii*. The external form of infected *P. clarkii* (A) had no difference with healthy ones, while the anatomical digestive tract (B) was lighter in color. A large number of brown spots in the hepatopancreas and white stuff in jejunum were observed (C). HP: hepatopancreas, IT: intestine.

**Fig. 2.**
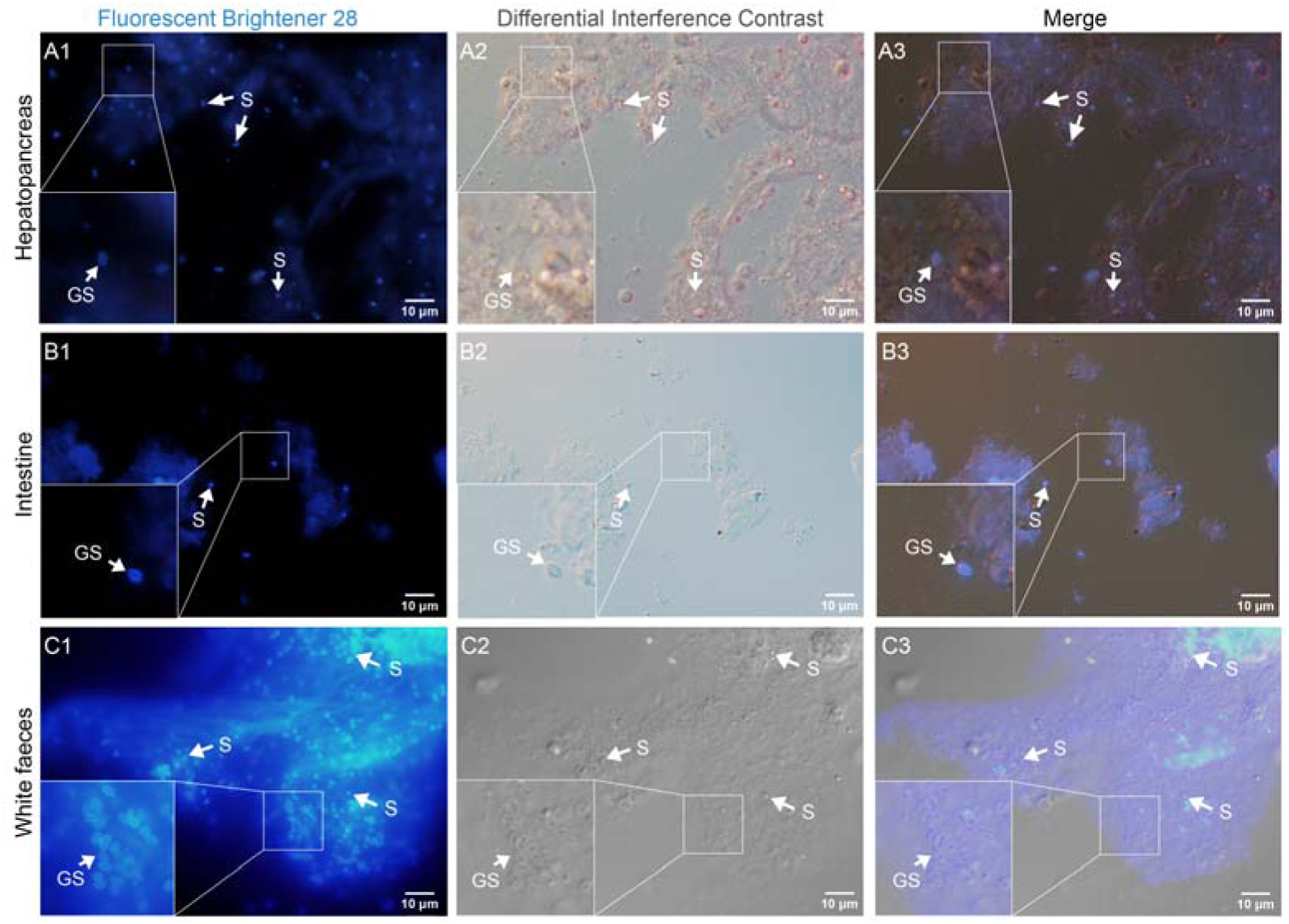
Fluorescence microscopic examination of hepatopancreas (A1-A3), intestine (B1-B3) and white feces (C1-C3) homogenate. Fluorescent Brightener 28 was used to stain with chitin layer of microsporidian spores. S: EHP mature spores; GS: germinated spore.

**Fig. 3.**
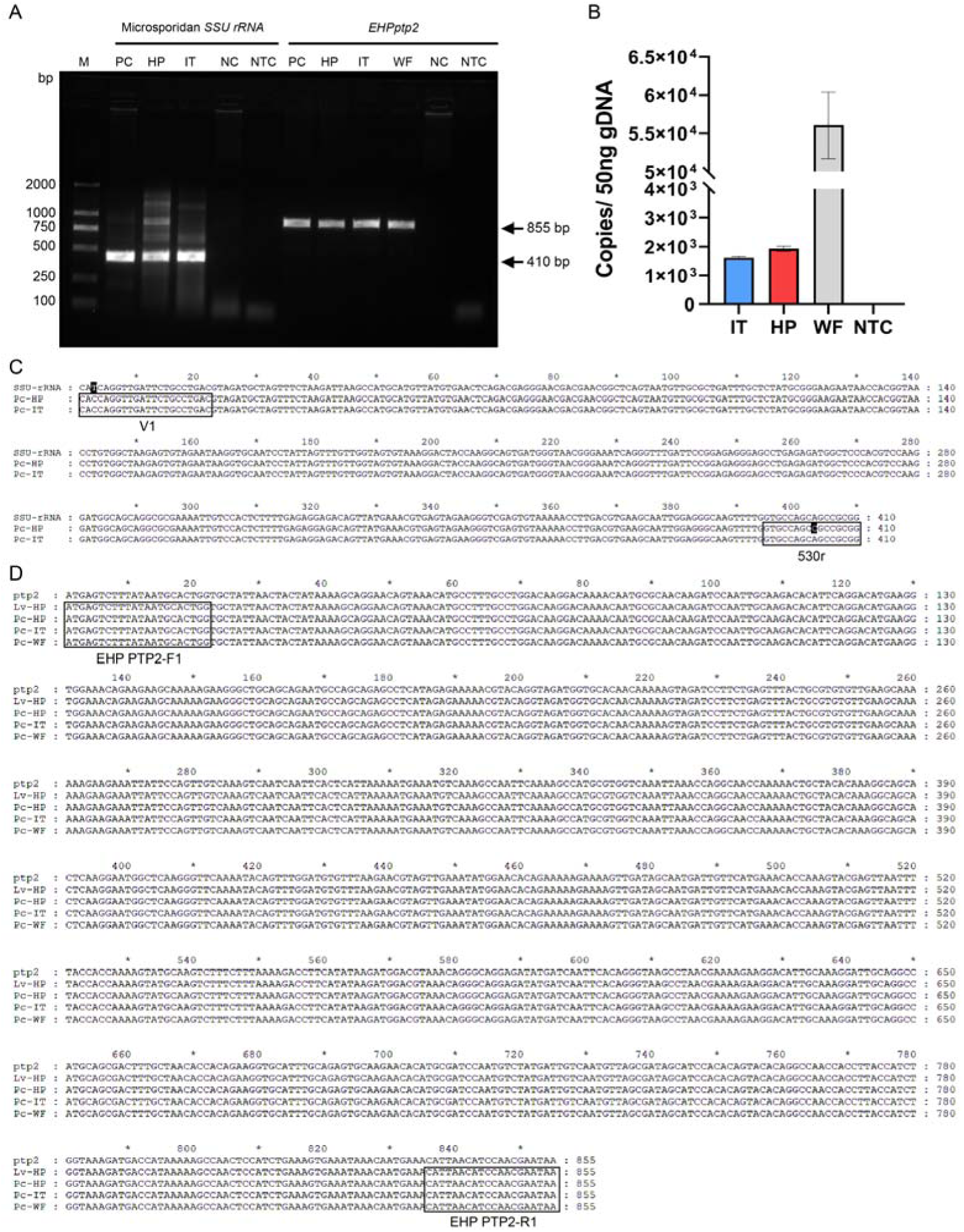
Certification of EHP loaded in the crayfish *P. clarkii*. A. PCR products amplified from tissues of infected *P. clarkii*. B. Quantification of EHP in tissues of *P. clarkii* by *EHPptp2* qPCR. Sequence alignments of *SSU rRNA* (C) and *EHPptp2* (D) amplicons and reference. PC: positive control, PCR products amplified from EHP infected *L. vannamei*; NC: negative control, no products were amplified from EHP-free *P. clarkii*; NTC: no template control, ddH_2_O was used as template; SSU-rRNA: small subunit ribosomal RNA (GenBank: MNPJ01000021.1); ptp2: polar tube protein 2 of EHP (GenBank: No. MT249228.1); Pc: *Procambarus clarkii*; Lv: *Litopenaeus vannamei*; HP: hepatopancreas; IT: intestine; WF: white faces.

### 3.2 Histological examination showed tissue damage in hepatopancreas and intestine of EHP infected *P. clarkii*

H&E stained paraffin section showed a various degree of tissue lesion in the EHP-positive *P. clarkii* (Fig. 4). The hepatopancreas showed severely necrotic pathologic-changes that the structure of hepatopancreatic tubules was damaged and infiltrated by a large amount of hemolymph accompanied by a severe hemocyte aggregation (Fig. 4 A1). The basal layers of hepatopancreatic tubules were ruptured along with epithelial cells disruption and shed, and EHP spores distributed among the cytoplasm of necrotic epithelial cells and lumen of necrotic tubules (Fig. 4 A1-A3 & C1-C3). While in healthy tissues, the hepatopancreatic tubules maintain intact morphological structure with epithelial cells tightly bound to the basal membrane, and the hemocyte was blocked from hepatopancreatic tubule epithelial cells (Fig. 4 B & D).

**Fig. 4.**
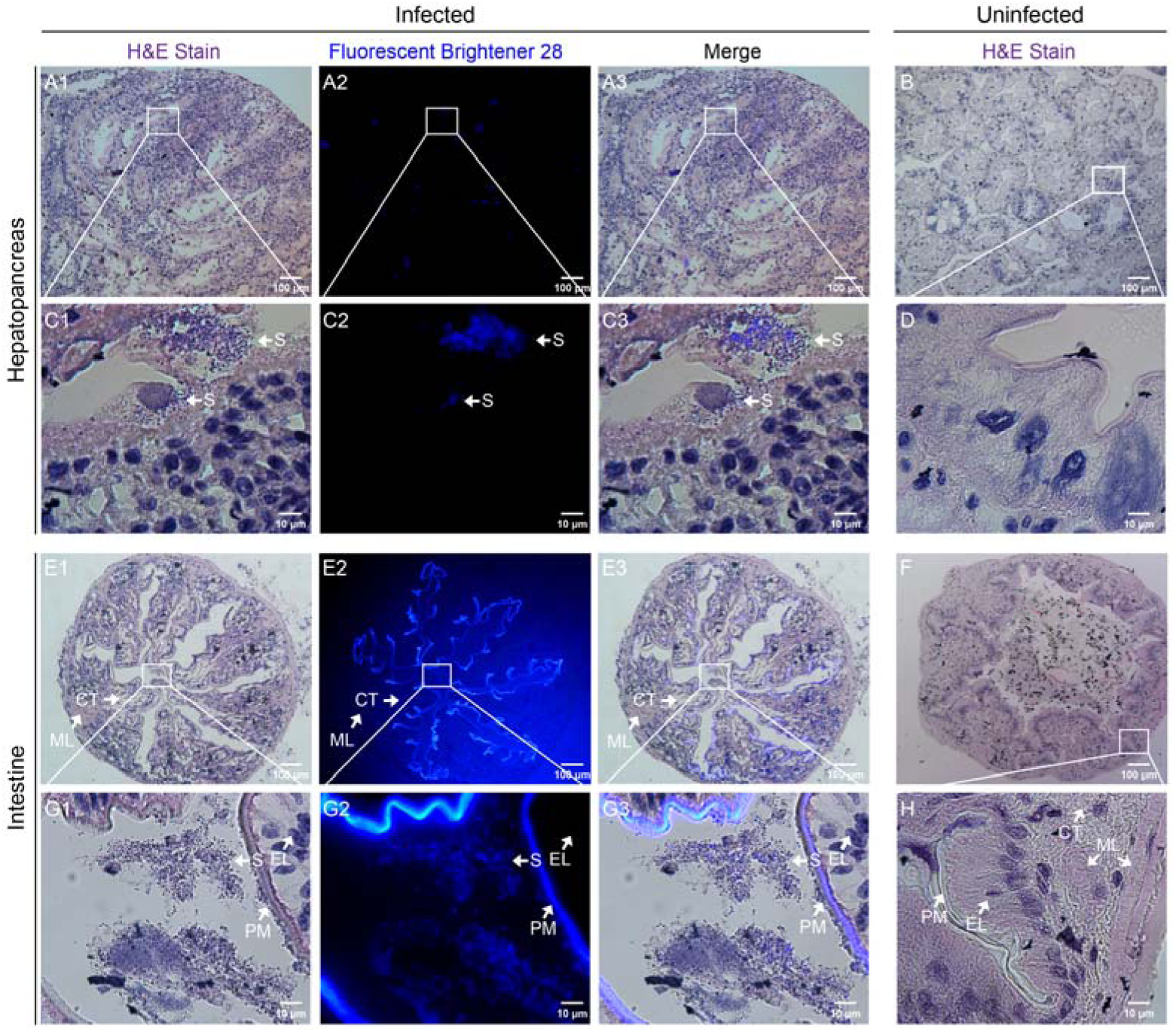
H&E-stain showed the tissue damage in the EHP infected *P. clarkii*. Abundant necrotic areas were seen in the infected hepatopancreas tissue and the morphology of hepatopancreatic tubules were difficult to distinguish (A1-A3). Clustered basophilic EHP spores were distributed among the necrotic areas (C1-C3) compared with those of EHP-free ones (B & D). The intestine of infected crayfish exhibited distinct epitheliolysis, enlarged interstices and sharped pleated ridges (E1-E3 & G1-G3) compared with those of EHP-free ones (F & H). S: EHP mature spores; PM: peritrophic membrane; EL: epithelial layer; CT: connective tissue layer; ML: muscle layer.

The intestine exhibited abnormal characteristics that epithelial cells necrosis, cytoplasmic staining was not obvious, the interstices in the tissue increased, the edges of the pleated ridge were sharper, and the muscle layer atrophied slightly (Fig. 4 E1-E3, G1-G3). In EHP-free crayfish, the peritrophic membrane, epithelial layer, connective tissue layer and muscle layer were tightly ordered (Fig. 4 F & H). There were some inclusions in the lumen (Fig. 4 F).

### 3.3 Developing stages of EHP were found in the hepatopancreatic cells of *P. clarkii*

In order to specifically indicate different stages of EHP, the polyclonal antiserum against EHP proteins extracts was prepared. Anti-EHP serum can recognize developing stages and mature spores of EHP. Combined with specific anti-EHP serum and Fluorescent brightener 28 labeling chitin layer of spores, proliferative stage, sporogonic stage and mature spores can be distinguished (Chen et al., 2017; Senderskiy et al., 2021). Early merogonic plasmodia, characterized by what anti-EHP serum can recognize without obvious chitin layer signal, were found in the hepatopancreas of EHP infected *P. clarkii* (Fig. 5 A1-C5). Sporogonic stages and mature spores were abundant in the hepatopancreas and intestine (Fig. 5 & Fig. S3). No hybridization signal was observed in EHP infected *P. clarkii* slices with unimmunized antiserum (Fig. S3) and in EHP-free *P. clarkii* slices with anti-EHP serum (Fig. S4).

**Fig. 5.**
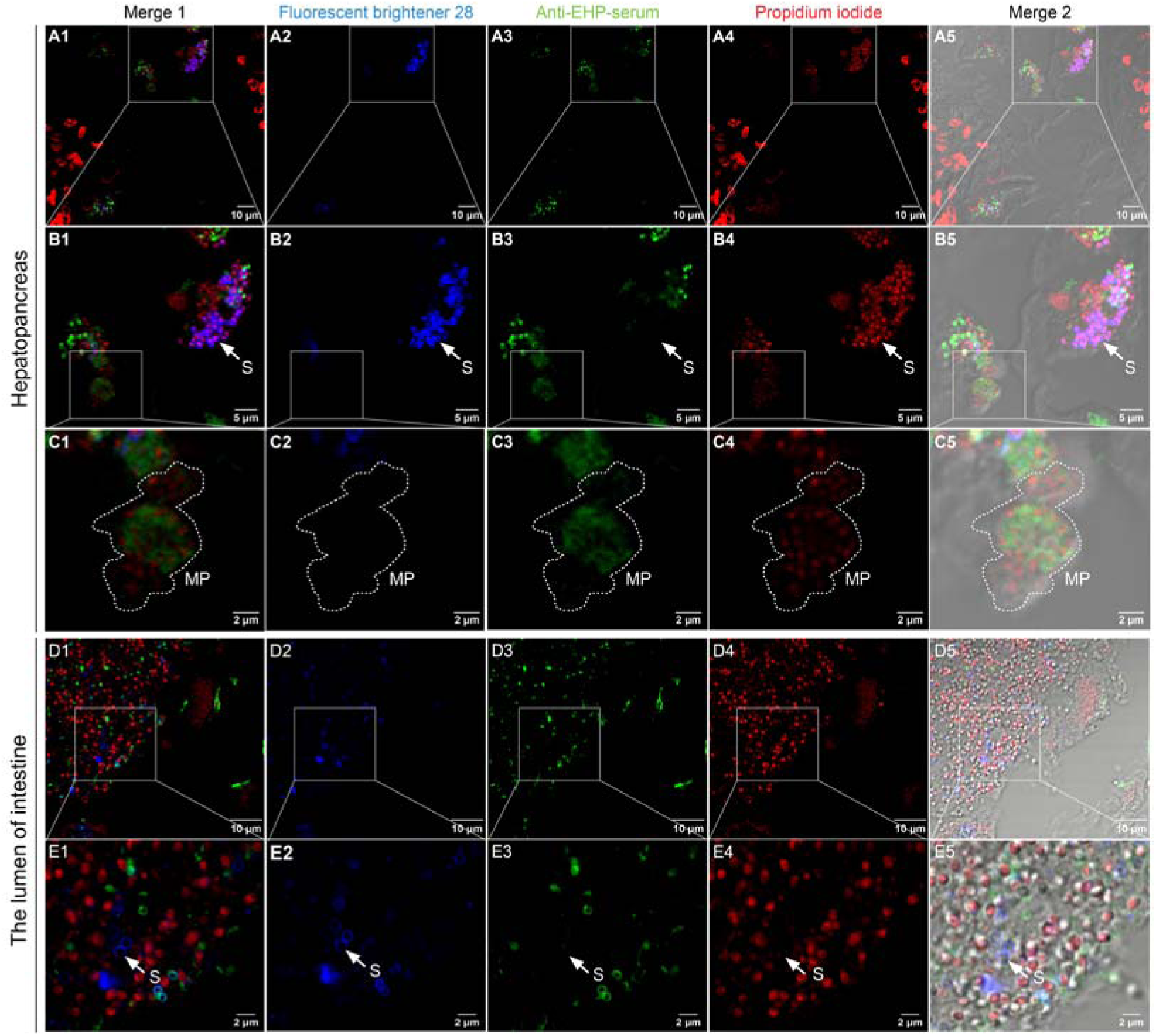
Developing stages of EHP in the hepatopancreas (A1-C5) and the lumen of intestine (D1-E5) of crayfish illustrated by indirect-immunofluorescent assay. EHP was labeled with anti-EHP serum (green), chitin coats of spores were labeled using Fluorescent brightener 28 (blue) and nuclei was stained by Propidium iodide (red). The regions selected by the rectangular boxes were magnified to the image below. Mature spores (B1-B5) and proliferative stage (C1-C5) of EHP can be seen in the hepatopancreas of crayfish. Different stages of EHP mix in the lumen of infected intestine (D1-E5). MP: merogonic plasmodia; S: EHP mature spores.

TEM further demonstrated the proliferation of EHP in crayfish hepatopancreatic cells. Numerous mature spores and sporogonic stages of EHP clustered in the cytoplasm (Fig. 6 A & B). Some sporogonic spores distributed near the host lipid droplet, and the membrane of lipid droplet contacted closely by the spores was deformed (Fig. 6 A & C). The mature spores were clearly characterized by exospore, endospore, plasm membrane, polaroplast, anchoring disc, single nucleus and 5-6 coils of polar filament (Fig. 6 E & F). EHP in sporogonial phase (during the sporogonic stage) was distinguished by thinner chitin layer and more unordered polar filaments (Fig. 6 E). Some spores were surrounded by a membranous structure individually or in pairs (Fig. 6 B & D). The germination behavior in hepatopancreas was verified by a germinated polar tube connected with EHP spore (Fig. 6 F).

**Fig. 6.**
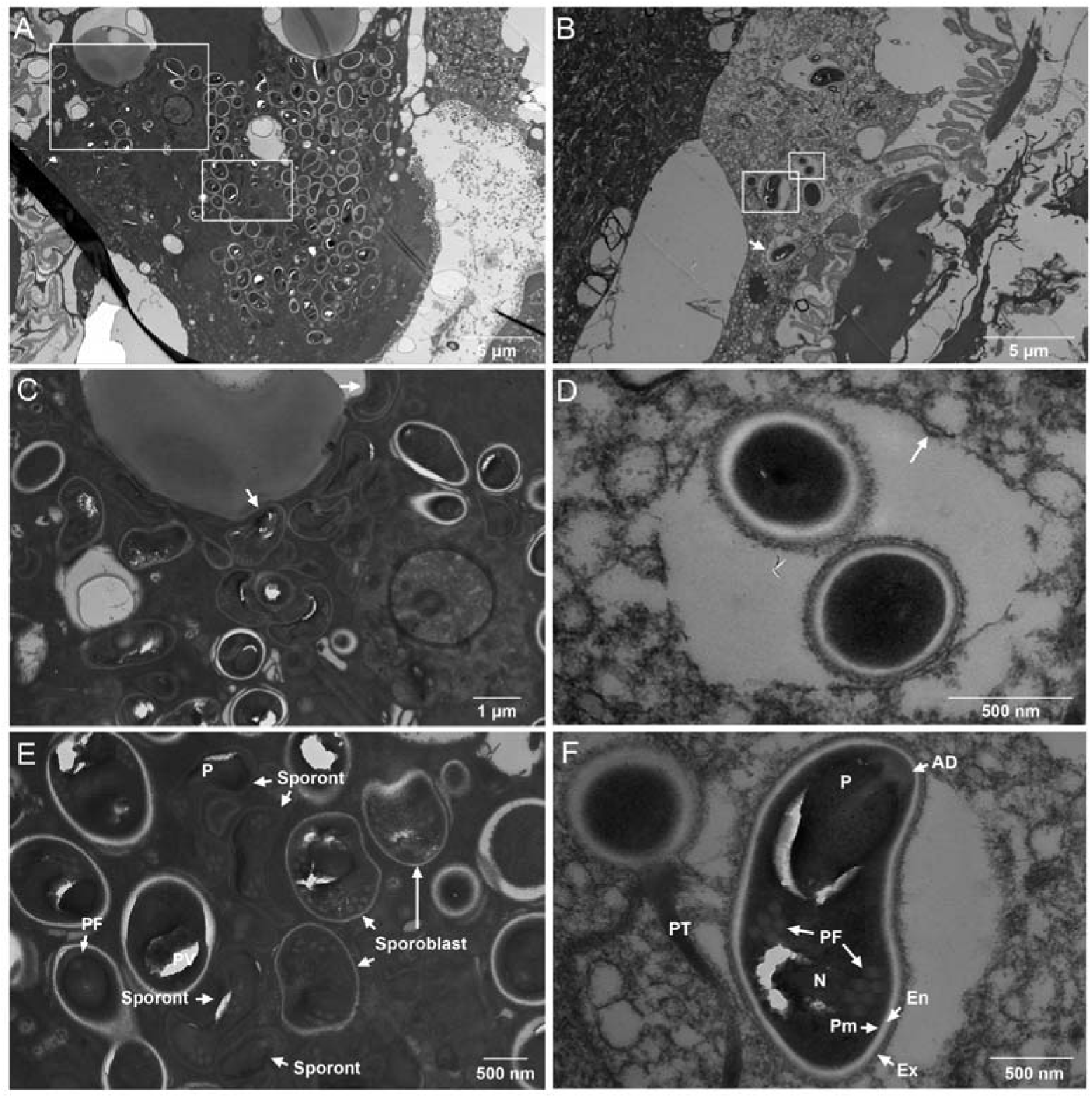
Ultrastructure of sporogonic stages and mature spores of EHP in hepatopancreatic epithelia of crayfish. Abundant sporogonic and mature stage of EHP were distributed in the cytoplasm of hepatopancreatic epithelial cells. Sporogonic spores were found contacted closely to the membrane of host lipid droplet, and the membrane was deformed (A & C). Several spores were enveloped by a membranous structure individually or in pairs (B & D). Mature spores were characterized by microsporidia ultrastructure with 5-6 coils of polar filament (F), while a thin spore wall was in the sporogonic phase of EHP (E). Image C and E, D and F were the magnification of the regions selected by the rectangular boxes in image A and B respectively. Ex: exospore; En: endospore; Pm: plasm membrane; P: polaroplast; PF: polar filament; PT: polar tube; N: nucleus; AD: anchoring disc.

## 4. Discussion

*L. vannamei* of and *P. clarkii* cultivation are the two most productive aquatic animals. The world production of their exceeds 8 million tons, accounting for more than 70% of the farmed crustaceans (FAO, 2022). With the rapid expansion of breeding scale, bacteria, viruses, parasites and other diseases have brought great trouble for aquaculture. It is noteworthy that the effects of pathogens vary from periods and species. For instance, the impact of White spot syndrome virus (WSSV) on crayfish was not obvious at first (Chou et al., 1995), but now it becomes one of the most threatening disease agents (Baumgartner et al., 2009; Dragicevic et al., 2021). EHP was initially neglected due to the prevalence of acute hepatopancreatic necrosis disease (AHPND) and White spot syndrome (WSD), while it is one of the most influential pathogen in shrimp farming now (Tang et al., 2015). In 2022, EHP is classified as a pathogen of Class II animal diseases in China. The pathogen found to proliferate in *P. clarkii* is noteworthy and alarming.

Microsporidia are important ecological regulators in nature. When the population of a species is massive increase, microsporidia will then proliferate and play a role in population size regulation. Compared with viral and bacterial diseases, EHP is characterized by relatively low mortality and huge economic loss (Patil et al., 2021). Therefore, it is particularly important to prevent intra- and interspecies infection of EHP to reduce the risk at the source.

The color of hepatopancreas and intestine turned white, which is a signal of EHP infection. This phenomenon may be related to lipid reduction in the crab and shrimp (Ding, 2021; Wu et al., 2022), and the presence of a large number of spores in the *P. clarkii* (Fig. 1 B & C; Fig. 2). EHP may supply its own development by absorbing and converting lipids from host. It is clear that EHP in the proliferative stage are closely bound to the lipid droplet of the host (Fig. 6 A & C). But there’s no way to see white hepatopancreas and intestine visually in *P. clarkii*. Because of the opaque exoskeleton, and EHP infected *P. clarkii* have no lesions on their appearance, it is hard to find crayfish infected. Flexible crawling ability also allows *P. clarkii* infected with EHP spread pathogens between different water bodies.

It is interesting that the crayfish in the same batch of samples (n=10) were negative to EHP by real-time qPCR after 2 weeks. However, the intestines of the same batch of crayfish were similar to that after starvation, with atrophy of the intestinal villi and enlargement of the intestinal lumen (Fig. 4 G). We suspect that probably due to regeneration of hepatopancreas epithelial cells in crayfish. The *P. clarkii* infected with EHP in the beginning, with daily water change, may somehow drained EHP out and recovered finally, since the regeneration of the complete hepatopancreas epithelium of crayfish seems to last less than 2 weeks (Davis and Burnett, 1964; Strus et al., 2019; Vogt, 2019). The short-term expulsion of large number of active EHP spores is beneficial for the transmission of EHP, but harmful to the farmed animals it parasitizes. However, this strategy in crayfish has implications for treatment of EHP infected shrimps.

In general, this article is a warning to the shrimp and crayfish aquaculture. Our results provide evidence that EHP can infect and proliferate in the hepatopancreas of crayfish *P. clarkii*. The crayfish prefer to swim upstream, resting in lakes, rivers, reservoirs, swampy ponds and ditches, and sometimes in the fields. Combining these features, *P. clarkii* was able to approach shrimp ponds and spread EHP they carried, or healthy crayfish entered EHP infected shrimp ponds, become infected accidentally and carry EHP to other healthy ponds or their habitats. All these remind farmers to take certain measures to avoid direct contact between *L. vannamei* and *P. clarkii* in culture processes. For example, it is forbidden to exchange ponds between the two species, to prevent wild *P. clarkii* from entering shrimp ponds.

## 5. Conclusion

The study showed EHP can proliferate in the hepatopancreatic cells of *P. clarkii*, and cause damage of hepatopancreas and intestine. Considering the large-scale cultivation of *L. vannamei* and *P. clarkii*, overlap farming areas between these two species as well as the ability of crayfish to crawl on land and water, our finding indicates the potential role of *P. clarkii* in the transmission of EHP, and this alerts us to the potentially greater risk of EHP in crayfish and shrimp aquaculture.

## Abbreviation

Pc: Procambarus clarkii
Lv: Litopenaeus vannamei
EHP: Enterocytozoon hepatopenaei
AHPND: acute hepatopancreatic necrosis disease
WSD: White spot syndrome
HP: hepatopancreas
WF: white faces
IT: intestine
PM: peritrophic membrane
EL: epithelial layer
CT: connective tissue layer
ML: muscle layer
S: EHP mature spores
MP: merogonic plasmodia
Ex: exospore
En: endospore
Pm: plasm membrane or sporoplasm membrane
P: polaroplast
PF: polar filament
PT: polar tube
N: nucleus
AD: anchoring disc
EDD: electron-dense disk
EHPptp2: EHP polar tube protein 2
cox: mitochondrial cytochrome oxidase c subunit I
D-loop: mitochondrial control region (Displacement loop region)
gDNA: genomic DNA
IFA: indirect-immunofluorescence assay
TEM: transmission electron microscopy
HPM: hepatopancreas microsporidiosis
SPF: specific pathogen free
PC: positive control
NC: negative control;
NTC: no template control.

## Funding

This research was funded by the Natural Science Foundation of Chongqing: 2022YSZX-JCX0010CSTB and cstc2021jcyj-cxttX0005.

## Authors’ contribution

Bingxin Ling: Writing - original draft, Methodology, Investigation. Yujiao Wu: Writing - original draft, Methodology, and Validation. Qing Yu, Chunxia Wang and Mengjiao Hu: Investigation. Xianzhi Meng: Resources and Investigation. Mengxian Long: Conceptualization and Visualization. Guoqing Pan: Conceptualization, Writing - review & editing and Funding acquisition. Zhonghuai Xiang: Conceptualization, Funding acquisition. Zeyang Zhou: Supervision and Writing - review & editing. Jie Chen: Supervision, Writing - review & editing and Funding acquisition.

## Declaration of Competing Interest

The authors declare that they have no conflict of interest.

## Acknowledgments

The authors are very grateful to Qing Lv for the help of staining the TEM samples, and thank to Yuan Wang for revision of the manuscript.

## Notes

### Competing Interest Statement

The authors have declared no competing interest.

