## Supplementary materials for "*Enterocytozoon hepatopenaei* Proliferate in *Procambarus clarkii*: A Warning for Crayfish and Shrimp Aquaculture"

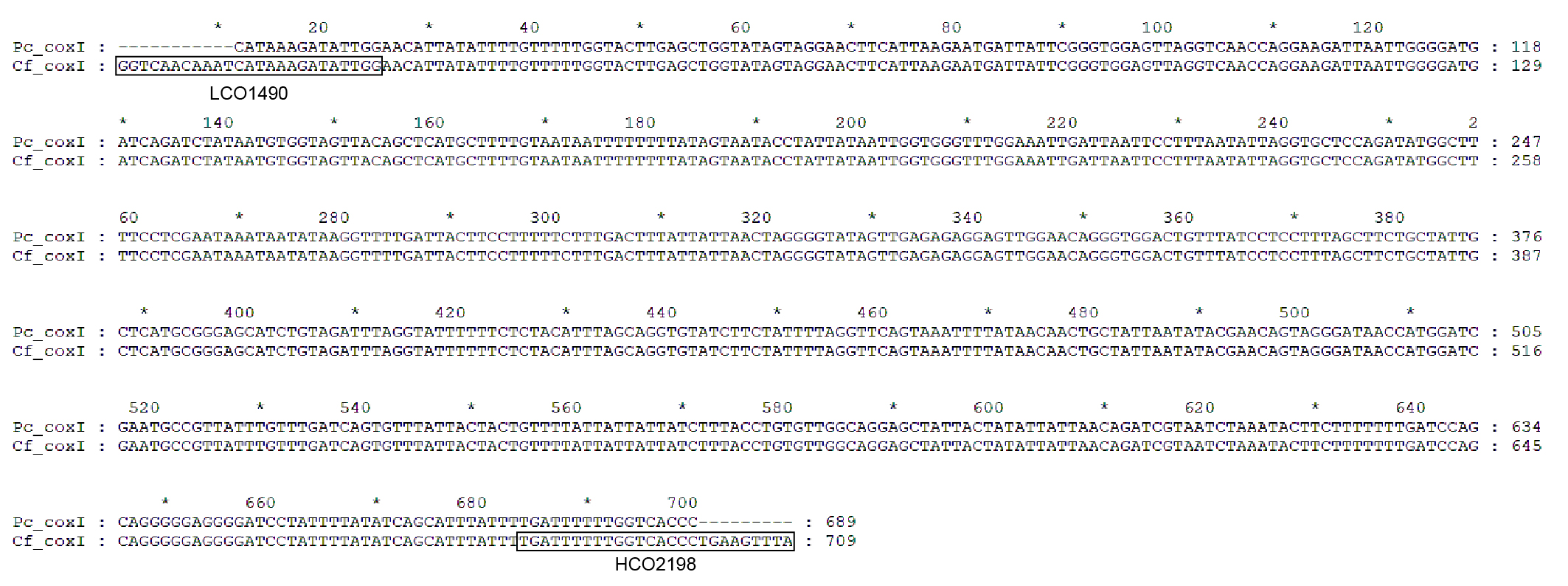

**FIGURE S1 Sequence alignment of cytochrome *c* oxidase subunitⅠ (*cox*Ⅰ) between *Procambarus clarkii* and sampled crayfish.** *Pc_cox*Ⅰ: *Procambarus clarkii* haplotype g cytochrome *c* oxidase subunit I gene, partial CDS of mitochondrial (GenBank: JN000903.1); *Cf_ coxⅠ*: sequenced *cox*Ⅰ of sampled crayfish.

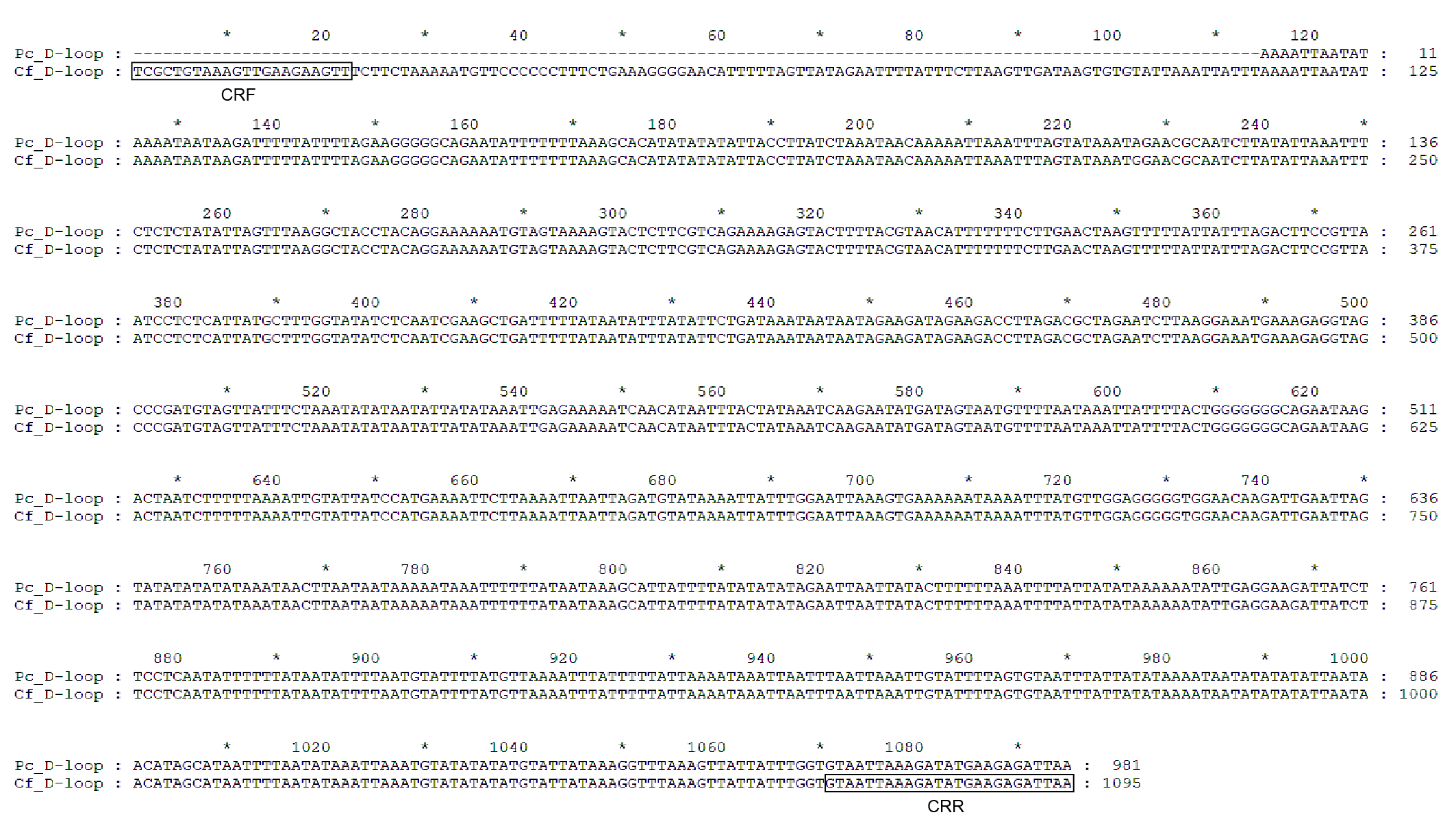

**FIGURE S2 Sequence alignment of mitochondrial control region (Displacement loop region, D-loop) between *Procambarus clarkii* and sampled crayfish.** *Pc_D-loop*: *Procambarus clarkii* haplotype d2 D-loop of mitochondrial, partial sequence (GeneBank: KC556829.1); *Cf_ D-loop*: sequenced D-loop of sampled crayfish.

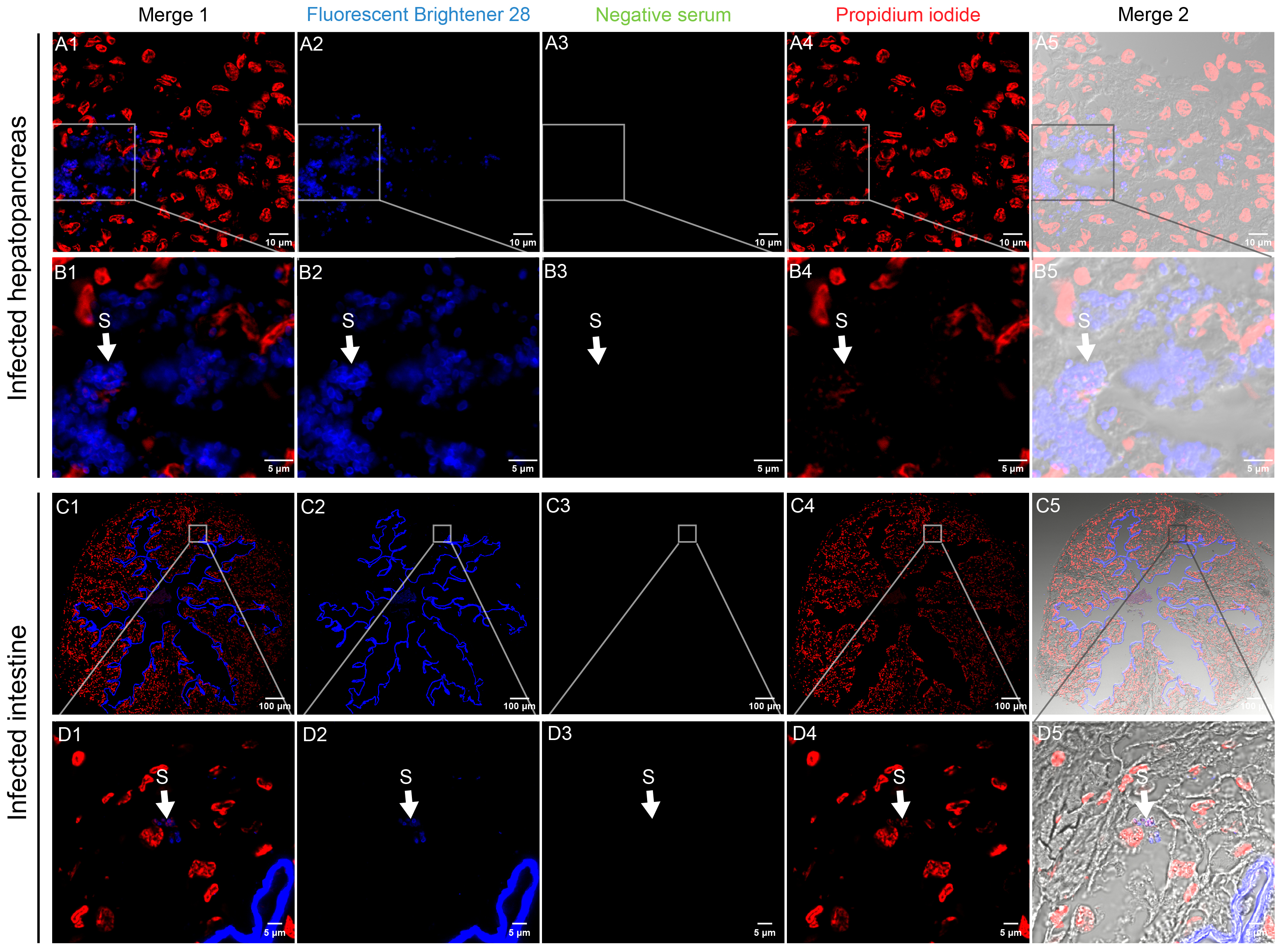

**FIGURE S3 Indirect immunofluorescence assay of infected *P. clarkii* with unimmunized antiserum (negative serum) showed no hybridization signal in the hepatopancreas (A1-B5) and intestinal tract (C1-D5).** Fluorescent brightener 28 (blue) and Propidium iodide (red) were used to stain chitin and nuclei respectively. Sporogonic stages and mature spores were characterized by overlap signal of chitin layer and nucleus. The regions selected by the rectangular box were enlarged to the image below. S: EHP mature spores.

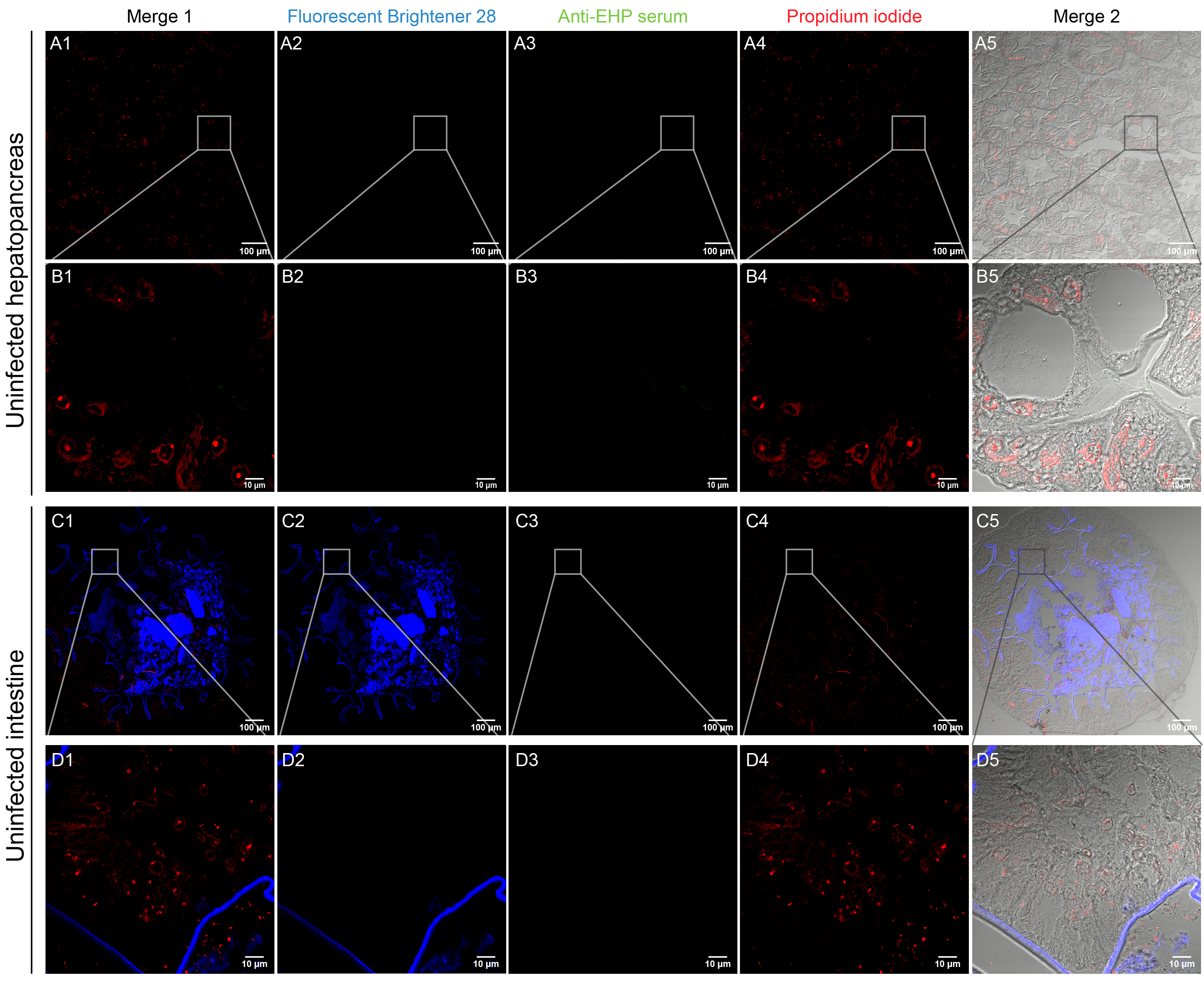

**FIGURE S4 Indirect immunofluorescence assay of uninfected *P. clarkii* with anti-EHP antiserum showed no hybridization signal in the hepatopancreas (A1-B5) and intestinal tract (C1-D5).** EHP were labeled with anti-EHP serum (green). Chitin layers of sporogonic stages and mature spores were labeled using Fluorescent brightener 28 (blue) and nuclei were stained by Propidium iodide (red). The regions selected by the rectangular box were enlarged to the image below.
